# All-Trans Retinoic Acid induces synaptic plasticity in human cortical neurons

**DOI:** 10.1101/2020.09.04.267104

**Authors:** Maximilian Lenz, Pia Kruse, Amelie Eichler, Julia Muellerleile, Jakob Straehle, Peter Jedlicka, Jürgen Beck, Thomas Deller, Andreas Vlachos

## Abstract

A defining feature of the brain is its ability to adapt structural and functional properties of synaptic contacts in an experience-dependent manner. In the human cortex direct experimental evidence for synaptic plasticity is currently missing. Here, we probed plasticity in human cortical slices using the vitamin A derivative all-trans retinoic acid, which has been suggested as medication for the treatment of neuropsychiatric disorders, e.g., Alzheimer’s disease. Our experiments demonstrate coordinated structural and functional changes of excitatory synapses of superficial (layer 2/3) pyramidal neurons in the presence of all-trans retinoic acid. This synaptic adaptation is accompanied by ultrastructural remodeling of the calcium-storing spine apparatus organelle and requires mRNA-translation. We conclude that all-trans retinoic acid is a potent mediator of synaptic plasticity in the adult human cortex.

## INTRODUCTION

The ability of neurons to respond to specific stimuli with structural, functional and molecular changes, i.e., to express plasticity, plays a fundamental role in normal brain function (Citri and Malenka, 2008). During the past decades cellular and molecular mechanisms of synaptic plasticity have been extensively studied in various animal models (Ho et al., 2011). Currently, however, no direct experimental evidence for coordinated structural and functional synaptic plasticity exists in the adult human cortex (Mansvelder et al., 2019). It thus remains unclear whether human neurons adapt their structural and functional properties, as seen, for example, in the rodent brain.

Vitamin A (all-trans retinol) and its metabolites have recently been linked to physiological brain functions such as axonal sprouting, synaptic plasticity and modulation of cortical activity (Drager, 2006; Shearer et al., 2012). Specifically, all-trans retinoic acid (atRA), which is clinically used in dermatology and oncology (Dobrotkova et al., 2018; Hu et al., 2009), has been studied for its neuroprotective and plasticity-promoting effects in animal models (Chen et al., 2014; Koryakina et al., 2009). Indeed, studies have been initiated that evaluate the effects of atRA in patients with brain diseases associated with cognitive dysfunction, e.g., Alzheimer’s disease, Fragile X syndrome, and depression (Bremner et al., 2012; Ding et al., 2008; Zhang et al., 2018). However, no direct experimental evidence exists for atRA-effects on neurons in the adult human cortex. Here, we assessed synaptic changes in atRA-treated human cortical slices prepared from neurosurgical resections, and we tested for the role of the plasticity-related protein synaptopodin, which is a key regulator of synaptic plasticity in the rodent brain (Deller et al., 2003; Segal et al., 2010; Vlachos et al., 2013) and has recently been linked to cognitive trajectory in human aging (Wingo et al., 2019).

## RESULTS

### All-trans retinoic acid treatment of human cortical slices

Cortical access tissue samples from 7 individuals who underwent clinically indicated neurosurgical procedures, e.g., for tumors or epilepsy, were experimentally assessed in this study (details provided in supplemental Table S1). Acute cortical slices (Gidon et al., 2020; Ting et al., 2018) were treated for 6 - 10 h with atRA (1 μM) or vehicle-only, and superficial (layer 2/3) pyramidal neurons were recorded in whole-cell configuration (Fig. 1A, B). While no significant differences in basic functional properties were detected between the two groups (Fig. 1C, D, E), a robust increase in the amplitudes of glutamate receptor-, i.e., α-amino-3-hydroxy-5-methyl-4-isoxazolepropionic acid (AMPA) receptor-mediated spontaneous excitatory postsynaptic currents (sEPSCs) was observed in the atRA-treated slices (Fig. 1F, G; see also Fig. S1). Mean sEPSC frequency was not significantly different between the two groups (Fig. 1G). These results are consistent with an atRA-mediated strengthening of excitatory neurotransmission onto human cortical pyramidal neurons.

**Figure 1.**
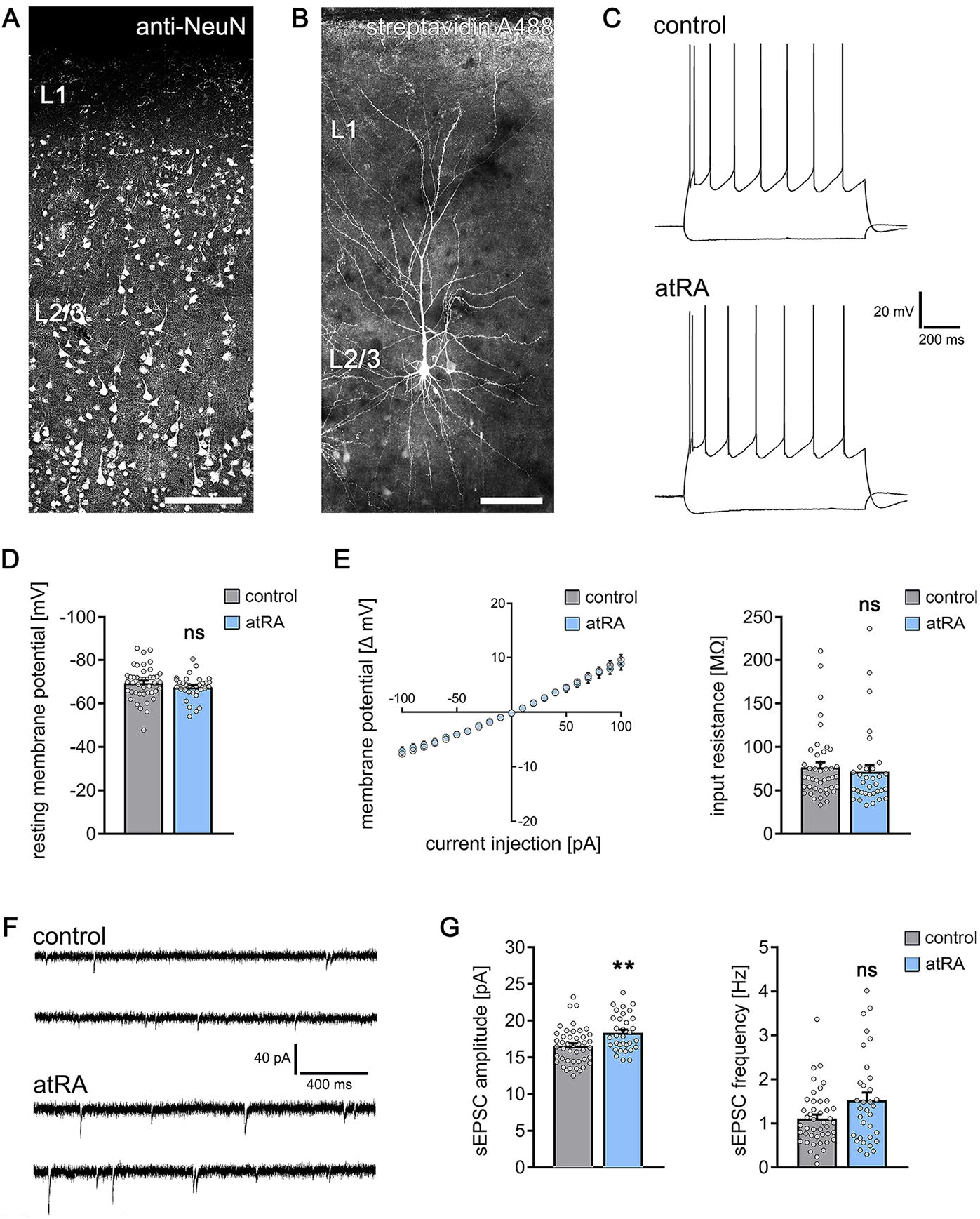
All-trans Retinoic Acid (atRA) induces plasticity of excitatory synapses in human cortical slices. (**A**) A representative human cortical slice stained for NeuN. Scale bar = 200 μm. (**B**) Recorded and post-hoc-labeled superficial (layer 2/3) pyramidal neurons. Scale bar = 100 μm. **(C-E)** Sample traces and group data of intrinsic cellular properties of cortical neurons from atRA- (1 μM, 6-10 h) and vehicle-only treated slices (responses to −100 pA and +350 pA current injection illustrated; n_control_ = 43 cells, n_atRA_ = 33 cells in 6 samples each; Mann-Whitney test). **(F, G)** Single cell recordings of AMPA receptor-mediated spontaneous excitatory postsynaptic currents (sEPSCs; n_control_ = 44 cells, n_atRA_ = 33 cells in 6 samples each; Mann-Whitney test, U = 442 for sEPSC amplitude analysis, p = 0.12 for sEPSC frequency; one cell in the atRA group with sEPSC amplitude = 38.4 pA and sEPSC frequency = 6.5 Hz was excluded from the analysis). Individual data points are indicated by gray dots. Values represent mean ± s.e.m. (ns, non-significant difference, ** p < 0.01).

### All-trans retinoic acid and dendritic spine morphology

A positive correlation between excitatory synaptic strength and dendritic spine sizes has been demonstrated in various animal models (Bosch and Hayashi, 2012; Matsuzaki et al., 2004). Hence, we wondered whether atRA induces structural changes of dendritic spines in cortical slices prepared from the adult human brain. To address this question, a set of recorded neurons was filled with biocytin and post-hoc stained using AlexaDye-labeled streptavidin to visualize dendritic morphologies (Fig. 2A). No significant differences in spine densities were observed between the two groups in these experiments (Fig. 2B). However, a marked increase in spine head sizes was evident in the atRA-treated group (Fig. 2C; c.f., Fig. 1). These findings establish a positive correlation between excitatory synaptic strength, i.e., sEPSC amplitudes, and dendritic spine sizes in human cortical pyramidal cells, thus revealing coordinated structural and functional changes of excitatory postsynapses in atRA-treated human cortical slices.

**Figure 2.**
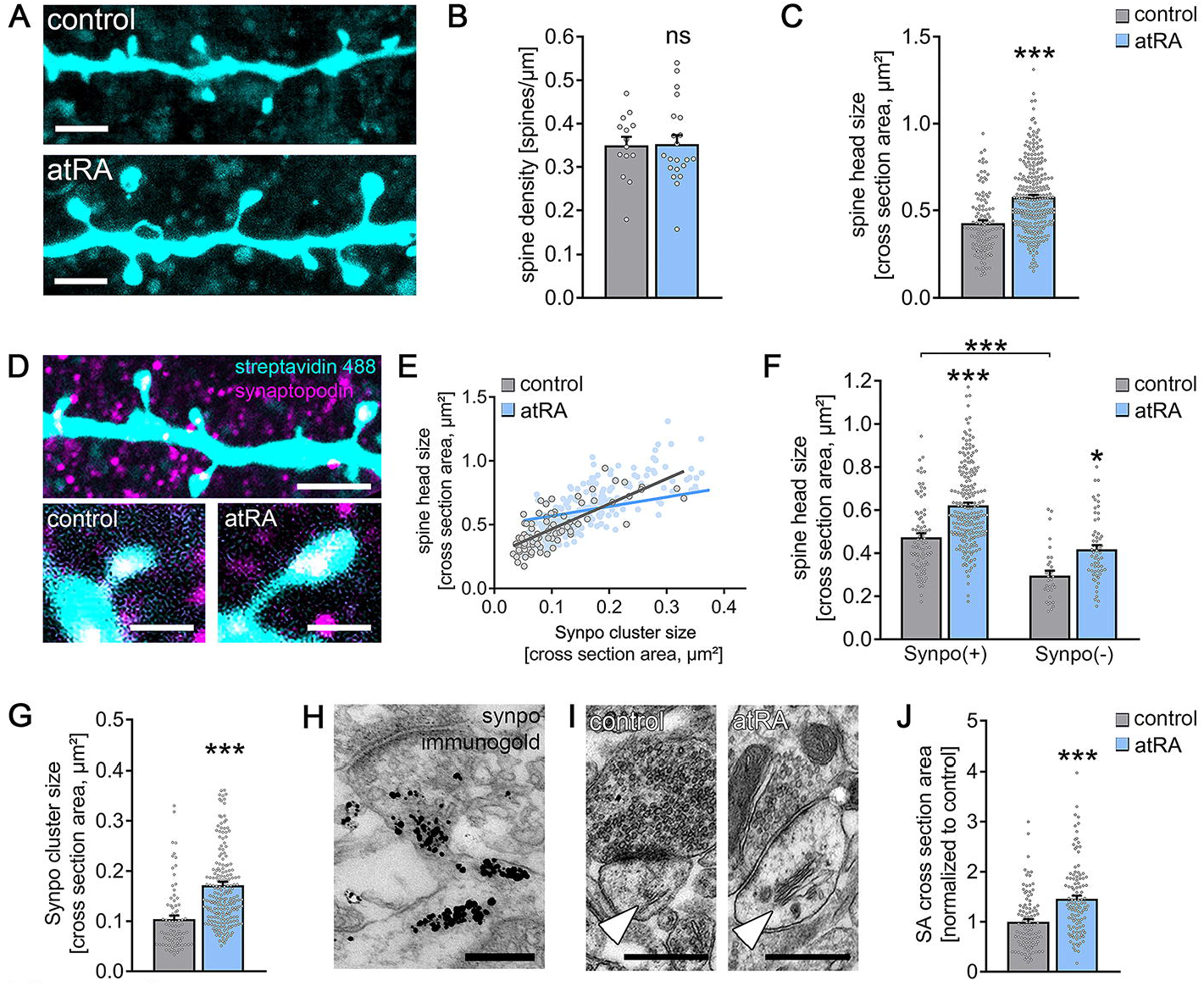
All-trans Retinoic Acid (atRA) induces dendritic spine plasticity in human cortical slices. **(A)** Example of dendritic segments of post-hoc-labeled superficial (layer 2/3) pyramidal neurons in atRA- (1 μM, 6-10 h) and vehicle-only treated slices. Scale bar = 3 μm. **(B, C)** Group data for spine densities (n_control_ = 14 dendritic segments, n_atRA_ = 21 dendritic segments from 3 samples each; Mann-Whitney test) and spine head sizes (n_control_ = 115 dendritic spines from 14 dendritic segments, n_atRA_ = 267 in 21 dendritic segments, 3 samples each; Mann-Whitney test, U = 8653). **(D)** Representative images of synaptopodin (Synpo) stained dendritic segments. Scale bar (upper panel) = 5 μm, Scale bar (lower panels) = 1 μm. **(E-G)** Correlation of synaptopodin cluster sizes and spine head sizes (n_control_ = 84 dendritic spines, n_atRA_ = 208 dendritic spines; Spearman r control = 0.73*** and atRA = 1***; three data points outside the axis limits in the atRA-treated group), group data of spine head sizes in synaptopodin-positive and -negative spines (synpo-positive spines: n_control_ = 84, n_atRA_ = 208; synpo-negative spines: n_control_ = 31, n_atRA_ = 59; Kruskal-Wallis test followed by Dunn’s post-hoc correction; one data point outside the axis limits in the atRA-treated group of synaptopodin-positive spines), and synaptopodin cluster sizes in the two groups (n_control_ = 84, n_atRA_ = 208; Mann-Whitney test, U = 3992; three data points outside the axis limits in the atRA-treated group). **(H)** Electron micrograph of synaptopodin immunogold-labeled spine apparatus (SA). Scale bar = 250 nm. **(I, J)** Examples and group data of SA (white arrowheads) cross sectional areas in atRA- and vehicle-only treated slices (n_control_ = 103, n_atRA_ = 114 from 3 samples each; values were normalized to the mean cross section area in the vehicle-only-treated group; Mann-Whitney test, U = 3489). Scale bar = 500 nm. Individual data points are indicated by gray dots. Values represent mean ± s.e.m. (ns, non-significant difference, * p < 0.05, *** p < 0.001).

### All-trans retinoic acid, synaptopodin and the spine apparatus organelle

In previous work, we demonstrated that the actin-modulating molecule synaptopodin (Mundel et al., 1997) is an essential component of the spine apparatus organelle (Deller et al., 2003), which is an enigmatic cellular organelle composed of stacked smooth endoplasmic reticulum found in subsets of dendritic spines (Kulik et al., 2019; Spacek, 1985). Synaptopodin-deficient mice do not form spine apparatuses and show defects in synaptic plasticity and behavioral learning (Deller et al., 2003). In this context, we showed that synaptopodin promotes the accumulation of AMPA receptors at synaptic sites (Maggio and Vlachos, 2014; Vlachos et al., 2009). Because atRA and synaptopodin have been linked to AMPA receptor-mediated synaptic plasticity (Aoto et al., 2008; Arendt et al., 2015a; Poon and Chen, 2008; Vlachos et al., 2009), we wondered whether atRA mediates its effects via synaptopodin and the spine apparatus organelle (Fig. 2D).

Synaptopodin clusters were detected in a considerable number of dendritic spines of human superficial (layer 2/3) pyramidal neurons: 74 ± 2% of all dendritic spines contained a synaptopodin cluster in characteristic positions, i.e., in the base, neck or head of spines (Fig. 2D, H). The previously reported positive correlation between synaptopodin cluster sizes and spine head sizes (Holbro et al., 2009; Vlachos et al., 2009) was also observed in the human cortex (Fig. 2E). Finally, comparison of synaptopodin-positive and synaptopodin-negative spines demonstrated that synaptopodin-positive spines are significantly larger than their synaptopodin-negative neighbors (Fig. 2F). Hence, synaptopodin clusters are found in strategic positions in a subset of large dendritic spines of the human cortex.

Systematic assessment of synaptopodin-positive and synaptopodin-negative spines revealed that atRA does not act specifically on synaptopodin-containing spines (Fig. 2F): A significant increase in spine head sizes was observed in both synaptopodin-positive and -negative dendritic spines. Although we did not observe a significant difference in the number of synaptopodin positive spines between the two groups (control: 71 ± 4% in 14 dendritic segments; atRA: 77 ± 2% in 21 dendritic segments; p = 0.25, Mann-Whitney test), atRA caused a significant increase in synaptopodin cluster sizes (Fig. 2G; c.f., Fig. 2E). We conclude that remodeling of synaptopodin clusters accompanies atRA-mediated coordinated structural and functional changes of dendritic spines in human cortical slices.

In previous work, we showed that remodeling of synaptopodin clusters reflects plasticity-related ultrastructural changes of spine apparatus organelles (Vlachos et al., 2013). We therefore wondered whether atRA changes ultrastructural properties of the spine apparatus organelle in human cortical neurons (Fig. 2H-J). After confirming that synaptopodin is a marker of the human spine apparatus organelle using pre-embedding immunogold stainings (Fig. 2H), ultrastructural properties of the spine apparatus organelle were assessed in transmission electron micrographs. Spine apparatuses were assessed in 103 (control) and 114 (atRA) cross sections of asymmetric synapses from three independent samples (Fig. 2I, J). Indeed, we found a marked increase in cross section area, which is consistent with the atRA-induced increase in synaptopodin cluster sizes (Fig. 2J; positive correlation of spine apparatus cross sections and spine cross sections shown in Fig. S2).

Taken together, these results establish a link between synaptopodin and the spine apparatus organelle in the human cortex. They demonstrate that changes in structural and functional properties of dendritic spines are accompanied by ultrastructural changes of spine apparatus organelles. We conclude that atRA is a potent mediator of coordinated (ultra-)structural and functional synaptic changes in the adult human cortex.

### Effects of atRA in synaptopodin-deficient mice

To learn more about the relevance of synaptopodin and the spine apparatus organelle in atRA-mediated synaptic plasticity we prepared acute cortical slices of the medial prefrontal cortex (mPFC) of synaptopodin-deficient mice (*Synpo*^*d/d*^; (Deller et al., 2003)). As shown in Figure 3A, atRA did not change sEPSC properties in superficial (layer 2/3) pyramidal neurons in the mPFC of *Synpo*^*d/d*^ mice (Fig. 3A), while a significant increase in sEPSC amplitudes was detected in mPFC slices of *Synpo*^*WT/WT*^ mice (control: 17.6 ± 0.5 pA, atRA: 19.9 ± 0.7 pA; n_control_ = 30 cells, n_atRA_ = 21 cells from three independent rounds each; p < 0.05*, Mann-Whitney test, U = 192). We repeated these experiments in acute slices prepared from *Thy1-GFP/Synpo*^*T/−*^ mice crossed to *Synpo*^*d/d*^ mice (Vlachos et al., 2013), which express the coding sequence of synaptopodin tagged to the green fluorescent protein (GFP) under the control of the Thy1.2 promotor. In a previous study, we demonstrated that the transgenic expression of GFP/Synpo rescues the ability of *Synpo*^*d/d*^ neurons to form spine apparatus organelles and to express synaptic plasticity (Vlachos et al., 2013). Indeed, a significant increase in sEPSC amplitudes was observed in mPFC layer 2/3 pyramidal neurons in these experiments (Fig. 3B). Hence, the presence of synaptopodin is required for atRA-mediated synaptic plasticity.

**Figure 3.**
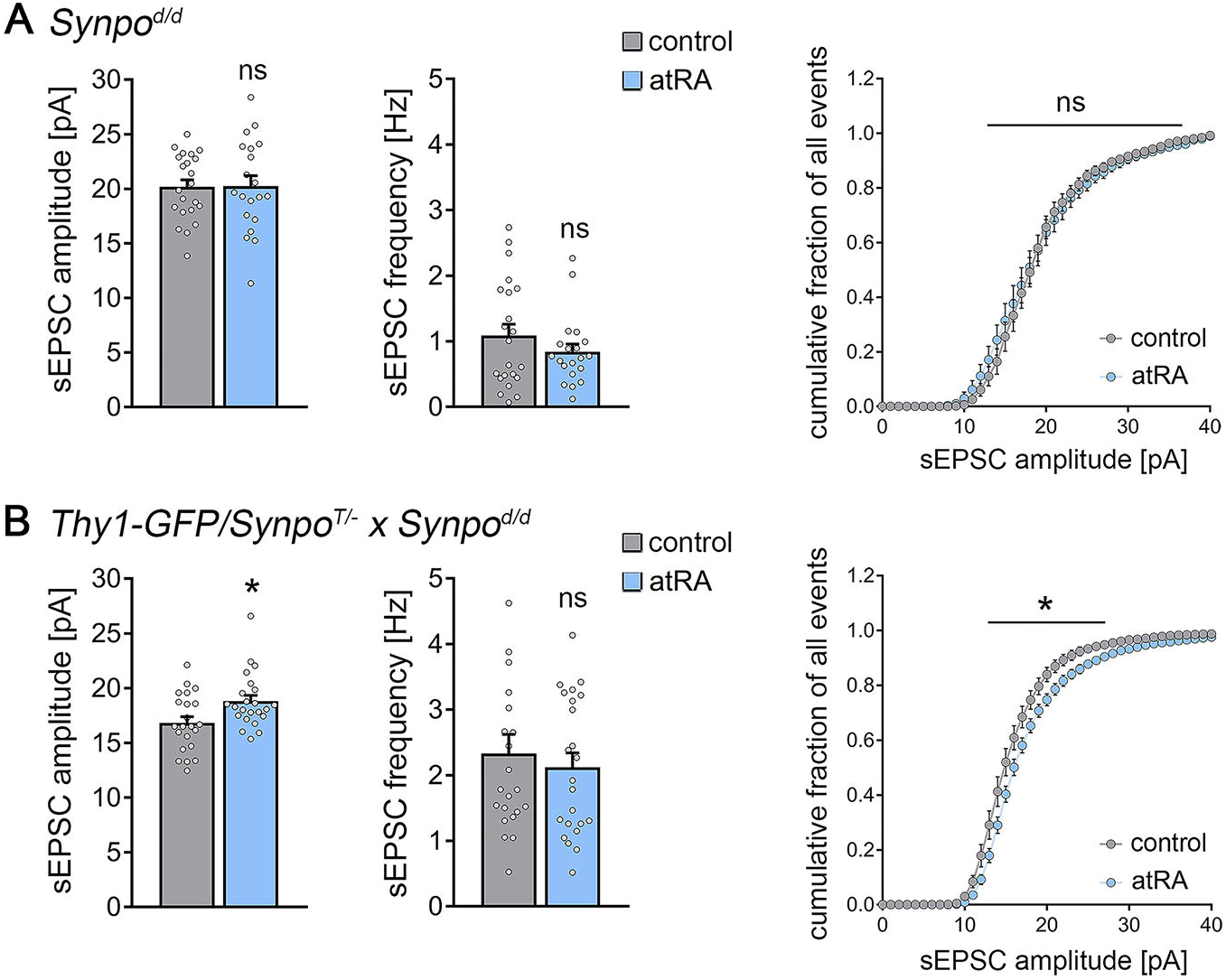
Effects of all-trans retinoic acid (atRA) in cortical slices prepared from synaptopodin-deficient mice. (**A**) Group data of AMPA receptor-mediated spontaneous excitatory postsynaptic currents (sEPSCs) recorded from superficial (layer 2/3) pyramidal neurons of the dorsomedial prefrontal cortex in slices prepared from synaptopodin-deficient mice (n_control_ = 22 cells, n_atRA_ = 20 cells in 3 independent experiments; Mann-Whitney test for column statistics and two-way ANOVA followed by Sidak’s multiple comparisons test for statistical evaluation of cumulative sEPSC amplitude distributions). (**B**) Group data of sEPSC recordings in cortical slices prepared from transgenic mice expressing GFP-tagged synaptopodin under the control of the Thy1.2 promotor crossed to synaptopodin-deficient mice (n_control_ = 22 cells, n_atRA_ = 23 cells in 3 independent experiments; Mann-Whitney test for column statistics, U_sEPSC amplitude_ = 157; two-way ANOVA followed by Sidak’s multiple comparisons test for statistical evaluation of cumulative sEPSC amplitude distributions; one data point outside the axis limits in sEPSC frequency control (vehicle-only) group). Individual data points are indicated by gray dots. Values represent mean ± s.e.m. (ns, non-significant difference, * p < 0.05).

### *In vivo* atRA treatment of synaptopodin-deficient mice

To provide further evidence for the role of synaptopodin in atRA-mediated synaptic plasticity, we resorted to *in vivo* long-term potentiation (LTP) of perforant path synapses in anesthetized mice (Fig. 4). atRA was injected intraperitoneally (10 mg/kg) and LTP was probed by electric stimulation of the perforant path with a theta burst stimulation protocol while recording evoked field excitatory postsynaptic potentials (fEPSPs) in the dentate gyrus of wildtype and *Synpo*^*d/d*^ mice (Fig 4A, B). As shown in Figure 4C, a significant increase in fEPSP slopes was observed in atRA-treated *Synpo*^*WT/WT*^ mice. However, atRA- and vehicle-only treated *Synpo*^*d/d*^ animals were indistinguishable in these experiments (Fig. 4D). These results confirm and extend our findings on the role of synaptopodin in atRA-mediated synaptic plasticity.

**Figure 4.**
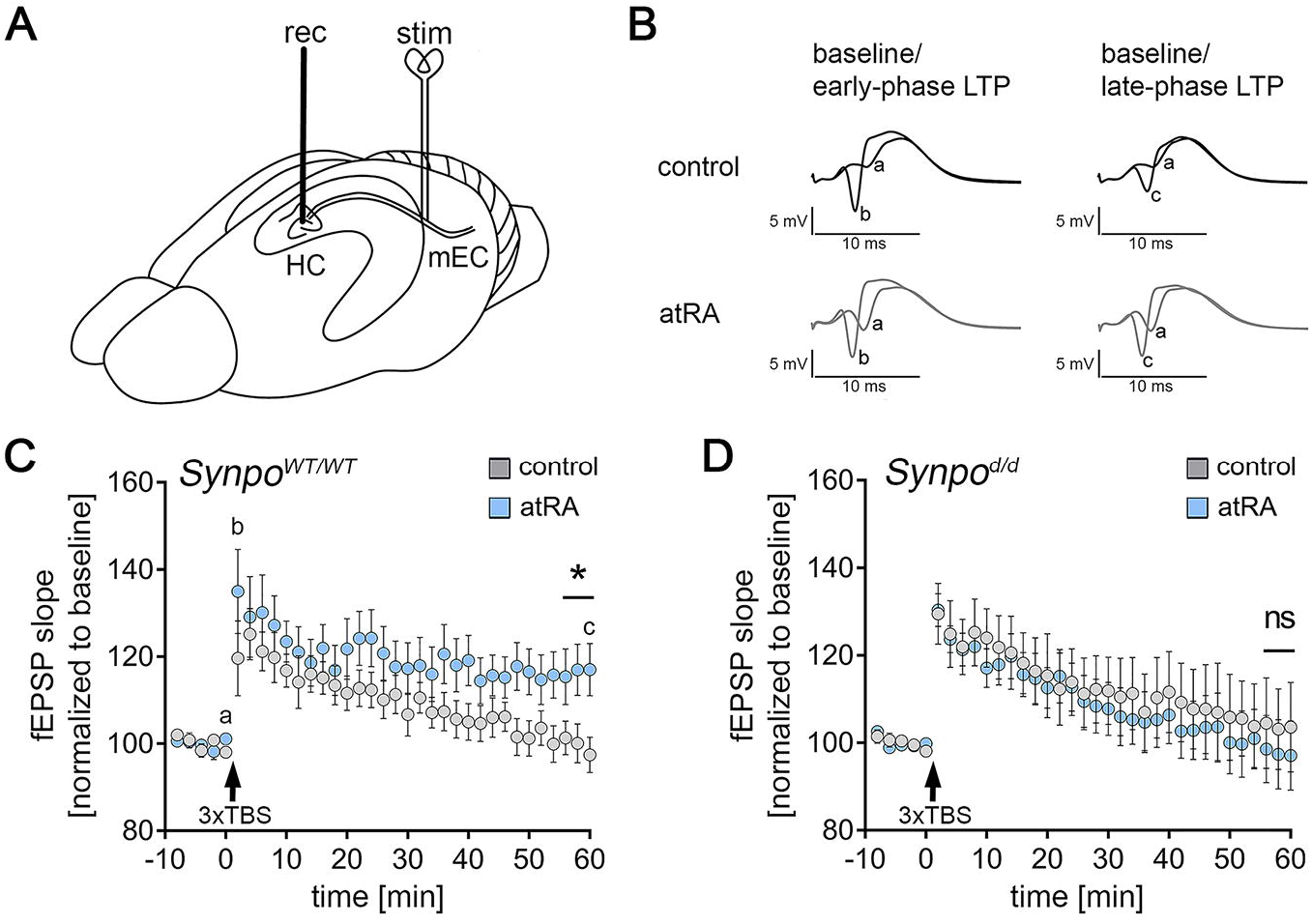
Intraperitoneal injection of all-trans retinoic acid (atRA) improves synaptic plasticity in wildtype but not synaptopodin-deficient mice. (**A**) Schematic illustration of experimental setting. The perforant path fibers, originating predominantly from the medial entorhinal cortex (mEC) are electrically stimulated while recording field excitatory postsynaptic potentials (fEPSP) from the dentate gyrus of the dorsal hippocampus (HC). (**B**) Representative traces of fEPSP recordings in wildtype mice at indicated points in time (a, b, c) after induction of long-term potentiation (LTP) in vehicle-only controls and atRA-injected mice (10 mg/kg, i.p., 3-6 hours prior to recordings) and vehicle-only-injected controls. **(C, D)** Group data of fEPSP slopes in the wildtype and synaptopodin-deficient mice (*Synpo*^*WT/WT*^: n_control_ = 9 animals, n_atRA_ = 9 animals, U = 13-17 for three terminal data points; *Synpo*^*d/d*^: n_control_ = 7 animals, n_atRA_ = 8 animals; Mann-Whitney test). Values represent mean ± s.e.m. (ns, non-significant difference, * p < 0.05).

### Pharmacologic inhibition of gene-transcription and mRNA-translation

atRA asserts its biological effects by acting at the gene-transcription and mRNA-translation level (Drager, 2006; Poon and Chen, 2008). Therefore, in a different set of acute cortical slices prepared from wildtype mice, actinomycin D (5 μg/ml) was used to block gene-transcription in the presence of atRA (1 μM). In these experiments, a significant increase in AMPA receptor-mediated sEPSC amplitudes was observed in the atRA group (Fig. 5A), thus confirming once more our major finding on atRA-mediated synaptic strengthening. We conclude from these results that major transcriptional changes do not explain the rapid effects of atRA on synaptic plasticity in our experimental setting. We next used anisomycin (10 μM) to block mRNA-translation, i.e., protein synthesis during the atRA treatment (Fig. 5B). Indeed, in the presence of anisomycin no significant changes in sEPSC amplitudes were detected, while sEPSC frequency was significantly reduced upon atRA treatment (Fig. 5B). These results suggest that atRA-mediated synaptic plasticity depends on mRNA-translation, i.e., protein synthesis.

**Figure 5.**
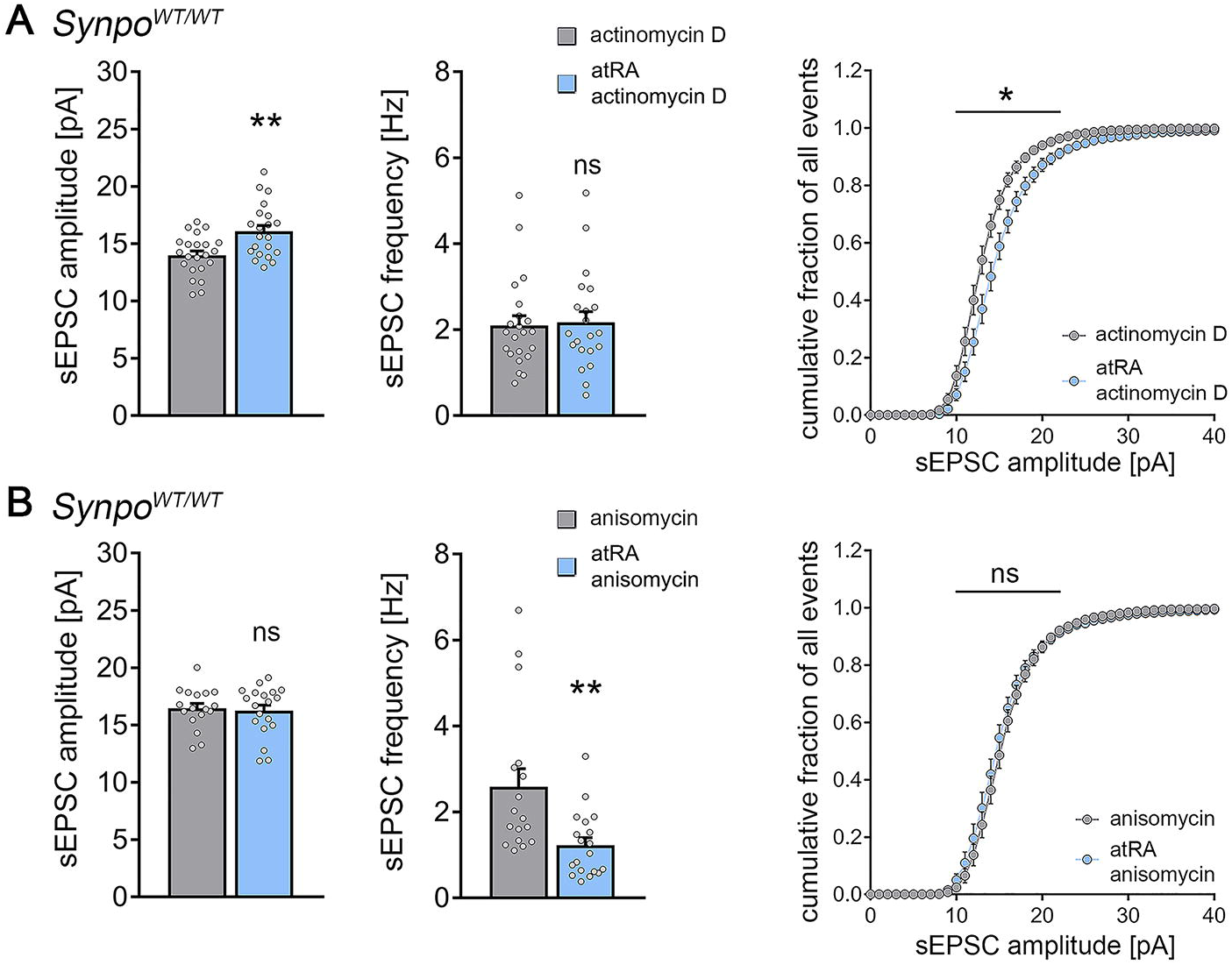
All-trans retinoic acid (atRA)-induced strengthening of excitatory synapses depends on mRNA-translation but not gene-transcription. (**A**) Group data of AMPA receptor-mediated spontaneous excitatory postsynaptic currents (sEPSCs) recorded from superficial (layer 2/3) pyramidal neurons of the dorsomedial prefrontal cortex in slices prepared from wildtype mice treated with actinomycin D (5 μg/ml; n_control_ = 22 cells, n_atRA_ = 21 cells in 3 independent experiments; Mann-Whitney test for column statistics, U_sEPSC amplitude_ = 120; two-way ANOVA followed by Sidak’s multiple comparison test for statistical evaluation of cumulative sEPSC amplitude distribution). (**B**) Group data of sEPSCs recordings from anisomycin-treated slices (10 μM; n_control_ = 17 cells, n_atRA_ = 19 cells in 3 independent experiments; Mann-Whitney test for column statistics, U_sEPSC frequency_ = 69; two-way ANOVA followed by Sidak’s multiple comparison test for statistical evaluation of cumulative sEPSC amplitude distribution). Individual data points are indicated by gray dots. Values represent mean ± s.e.m. (ns, non-significant difference, * p < 0.05, ** p < 0.01).

### Pharmacologic inhibition of mRNA-translation in human cortical slices

Based on these results obtained in the murine brain, we returned to human brain slices and tested whether mRNA-translation is also required for atRA-mediated structural and functional plasticity in the human cortex (Fig. 6). For this, acute cortical slices were prepared from human surgical access material. AMPA receptor-mediated sEPSCs were recorded from superficial (layer 2/3) pyramidal neurons of atRA- and vehicle-only-treated slices, where during the treatment period mRNA-translation was pharmacologically inhibited by anisomycin (10 μM). No significant differences in sEPSC amplitudes were observed between the two groups in these experiments (Fig. 6A). Consistent with these findings, anisomycin also blocked the atRA-mediated increase in spine head sizes and synaptopodin cluster sizes, while the previously reported difference in spine head sizes between synaptopodin-positive and synaptopodin-negative spines remained unchanged (Fig. 6B). Thus, atRA-mediated plasticity requires mRNA-translation, i.e., protein synthesis to trigger coordinated changes in synaptic strength, spine head sizes and synaptopodin cluster properties in the human cortex.

**Figure 6.**
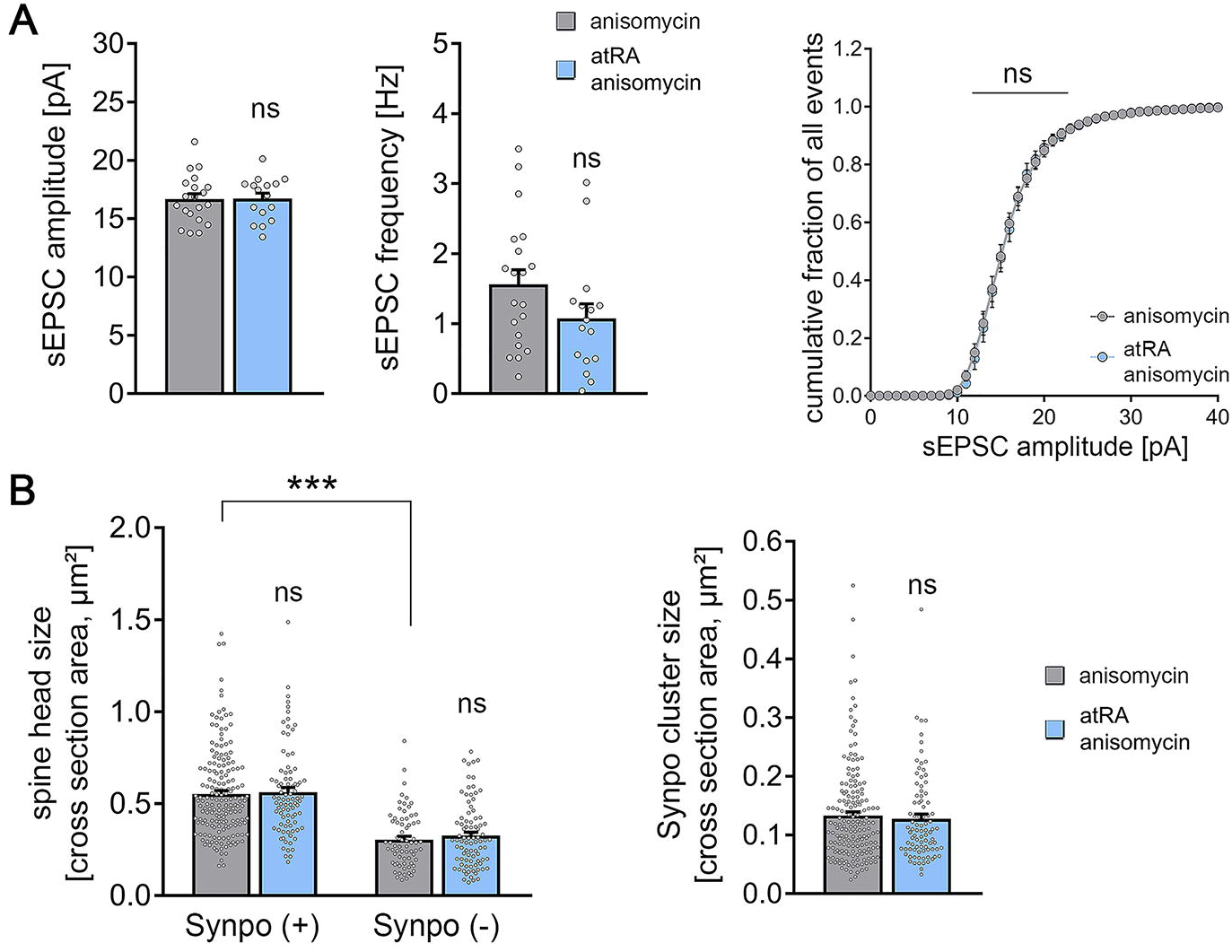
Pharmacologic inhibition of protein synthesis prevents all-trans retinoic acid (atRA)-induced synaptic plasticity in human cortical slices. (**A**) Group data of AMPA receptor-mediated spontaneous excitatory postsynaptic currents (sEPSCs) recorded from superficial (layer 2/3) pyramidal neurons. Human cortical slices were treated with atRA or vehicle-only in the presence of anisomycin (10 μM; n_control_ = 20 cells, n_atRA_ = 16 cells in 3 independent experiments; Mann-Whitney test for column statistics; two-way ANOVA followed by Sidak’s multiple comparison test for statistical evaluation of cumulative sEPSC amplitude distribution). (**B**) Spine head sizes and synaptopodin cluster sizes in vehicle-only and atRA (1 μM, 6-10 h) treated human cortical slices (c.f., Fig. 2 F, G; synaptopodin-positive spines: n_control_ = 175, n_atRA_ = 88; synaptopodin-negative spines: n_control_ = 69, n_atRA_ = 88, 1-11 segments per sample in 3 independent experiments; Kruskal-Wallis test followed by Dunn’s post-hoc correction for spine head size and Mann-Whitney test for synaptopodin cluster size). Individual data points are indicated by gray dots. Values represent mean ± s.e.m. (ns, non-significant difference, *** p < 0.001).

## DISCUSSION

Vitamin A and its metabolites regulate important biological processes by binding to nuclear and cytoplasmic receptors (Al Tanoury et al., 2013). They are key mediators of cell growth, cell survival and differentiation accounting for the teratogenic effects of atRA as well as its clinical use, e.g., in the therapy of promyelocyte leukemia (Hu et al., 2009). Studies in the central nervous system have mainly focused on embryonic and early postnatal development (Luo et al., 2004; Rataj-Baniowska et al., 2015). More recently, however, it has been recognized that retinoid metabolism and signaling occurs physiologically in the adult murine brain (Shearer et al., 2012). In this context, a series of animal studies revealed that retinoid receptors bound to specific mRNAs in the cytoplasm control synaptic plasticity by regulating synthesis, trafficking and accumulation of synaptic proteins (Aoto et al., 2008; Groth and Tsien, 2008; Maghsoodi et al., 2008; Poon and Chen, 2008). Some of these findings have recently been replicated in immature neurons derived from human inducible pluripotent stem cells (Zhang et al., 2018). The results of this study demonstrate the potential of atRA to induce plasticity in the adult human cortex. Consistent with recent reports on the relevance of protein synthesis in synaptic plasticity (Biever et al., 2020; Hafner et al., 2019; Sutton and Schuman, 2006) our experiments reveal that atRA-mediated synaptic plasticity in the adult human cortex requires mRNA-translation.

At the mechanistic level we identified synaptopodin as a key regulator of atRA-mediated synaptic plasticity. Our results demonstrate that synaptopodin is a marker of the human spine apparatus, which responds to plasticity-inducing stimuli with ultrastructural changes that accompany spine size changes. Both atRA and synaptopodin have been linked to the expression of Hebbian and homeostatic synaptic plasticity in rodent brain tissue and are associated with Ca^2+^-dependent AMPA receptors synthesis and trafficking (Arendt et al., 2015b; Vlachos et al., 2013; Vlachos et al., 2009). In turn, evidence exists that atRA and synaptopodin are linked to alterations of synaptic plasticity in pathological brain states (Bremner et al., 2012; Maggio and Vlachos, 2014). Notably, alterations in synaptopodin expression have been demonstrated in brain tissue of Alzheimer’s disease patients (Reddy et al., 2005) and a recent large-scale proteomic analysis study of the human brain associated synaptopodin with cognitive trajectory in the advanced age (Wingo et al., 2019). Considering that retinoids are discussed as medication for the treatment of Alzheimer’s disease-associated cognitive decline (Ding et al., 2008; Endres et al., 2014), it is conceivable that atRA may act – at least in part – by modulating synaptopodin expression, thereby recruiting the ability of neurons to express synaptic plasticity in the adult human cortex.

## Acknowledgments

We thank Barbara Joch and Sigrun Nestel for their excellent technical assistance.

## Funding

The work was supported by the EQUIP-Medical Scientist Program of the Faculty of Medicine, University of Freiburg to M.L., by the Berta-Ottenstein-Program for Clinician Scientists of the Faculty of Medicine, University of Freiburg to J.S., and by Deutsche Forschungsgemeinschaft (CRC 1080 to T.D. and A.V.; CRC 974 to A.V.).

## Author contributions

Roles of authors and contributors have been defined according to ICMJE guidelines and author contributions have been reported according to CRediT taxonomy. Conceptualization, M.L. and A.V.; Methodology, M.L., J.S., J.B. and A.V.; Formal Analysis, M.L., P.K., A.E. and J.M.; Investigation, M.L., P.K., A.E. and J.M.; Resources, J.B., T.D. and A.V.; Writing Original Draft, M.L., T.D. and A.V.; Visualization, M.L. and A.V.; Supervision, P.J. and A.V.; Project Administration, M.L., J.S., J.B. and A.V.; Funding Acquisition, M.L., J.S., T.D. and A.V.

## Competing interests

The authors declare no competing interests. T.D. received funding from Novartis for a lecture on human brain anatomy.

## Data and materials availability

All data is included in the main text or the supplementary materials. Original files are available upon reasonable request.

## METHODS

### Ethics statement

Human brain tissue was obtained from a local biobank operated in the Department for Neurosurgery at the Faculty of Medicine, University of Freiburg (AZ 472/15_160880). All experiments carried out in this study were performed based on positive evaluation by the local ethics committee (AZ 593/19) and/or according to the German animal welfare legislation and local authorities [approved by the animal welfare officers of the Faculty of Medicine at the University of Freiburg (AZ X-17/04C) and the Faculty of Medicine at the University of Frankfurt (AZ FU/1131)]. Animals were kept in a 12 h light/dark cycle with access to food and water ad libitum. Every effort was made to minimize distress and pain of animals.

### Preparation of acute human cortical slices

After resection, cortical access tissue was immediately transferred to oxygenated extracellular solution containing (in mM): 92 NMDG, 2.5 KCl, 1.25 NaH_2_PO_4_, 30 NaHCO_3_, 20 HEPES, 25 glucose, 2 thiourea, 5 Na-ascorbate, 3 Na-pyruvate, 0.5 CaCl_2_, and 10 MgSO_4_, pH = 7.3 to 7.4 at ~10°C [NMDG-aCSF; (Gidon et al., 2020; Ting et al., 2018)]. Prior to slicing procedure, cortical tissue was embedded in low melting agarose (Sigma Aldrich, #A9517; 1.8% (w/v) in phosphate buffered saline). 400 μm tissue sections were cut with a Leica VT1200S vibratome perpendicular to the pial surface in the same solution at 10°C under continuous oxygenation (5% CO_2_ / 95% O_2_). Slices were transferred to cell strainers with 40 μm pore size placed in NMDG-aCSF at 34°C and sodium levels were gradually increased as previously described (Ting et al., 2018). After recovery, slices were maintained for further experimental assessment at room temperature in extracellular solution containing (in mM): 92 NaCl, 2.5 KCl, 1.25 NaH_2_PO_4_, 30 NaHCO_3_, 20 HEPES, 25 glucose, 2 thiourea, 5 Na-ascorbate, 3 Na-pyruvate, 2 CaCl_2_, and 2 MgSO_4_. Cortical slices from all human samples looked macroscopically normal and showed no overt pathology.

### Preparation of acute mouse cortical slices

Adult mice (C57BL/6J, B6.129-Synpo^tm1Mndl^/Dllr (referred to as *Synpo*^*d/d*^) and B6.Cg-Synpo^tm1Mndl^Tg(Thy1-Synpo/GFP)1Dllr/Dllr (referred to as *Thy1-GFP/Synpo^T/−^ *x* Synpo^d/d^*); 6-11 weeks old) were anesthetized with isoflurane and rapidly decapitated. Brains were rapidly removed, washed in chilled (~10°C) aCSF and embedded in low-melting agarose (Sigma-Aldrich #A9517; 1.8% (w/v) in phosphate buffered saline). Coronal sections of the medial prefrontal cortex were prepared using a Leica VT1200S vibratome with the brain tilted dorsally at a 15° angle in NMDG-aCSF. Slice recovery and maintenance until experimental assessment as described above for acute human cortical slices.

### Pharmacology

Acute cortical slices prepared from one sample were randomly assigned to the respective treatment groups. All-trans retinoic acid (atRA) treatment was performed after recovering the acute slices by adding atRA (1 μM, Sigma Aldrich, #R2625) at a final concentration of 0.05% (v/v) DMSO to the holding extracellular solution. The control group from the same set of slices was treated equally but with vehicle-only (DMSO). Anisomycin (10 μM, Abcam, #ab120495) and actinomycin D (5 μg/ml, Sigma Aldrich, #A9415) were added to the holding solution 10 minutes before atRA was added. Sections were treated for at least 6 hours before experimental assessment.

### Whole-cell patch-clamp recordings

Whole-cell patch-clamp recordings of cortical pyramidal cells in the superficial layers (layer 2/3) were carried out at 35°C in a bath solution containing (in mM): 92 NaCl, 2.5 KCl, 1.25 NaH_2_PO_4_, 30 NaHCO_3_, 20 HEPES, 25 glucose, 2 thiourea, 5 Na-ascorbate, 3 Na-pyruvate, 2 CaCl_2_, and 2 MgSO_4_. In acute mouse brain slices, superficial (layer 2/3) pyramidal cells in the dorsomedial prefrontal cortex (anterior cingulate and prelimbic regions) were visually identified using a LN-Scope (Luigs & Neumann, Ratingen, Germany) equipped with an infrared dot-contrast and a 40x water-immersion objective (Olympus, NA 0.8). In human cortical slices, superficial (layer 2/3) pyramidal cells were visually identified at a distance range from 500 to 1000 μm from the pial surface on the pia/white matter-axis. Electrophysiological signals were amplified using a Multiclamp 700B amplifier, digitized with a Digidata 1550B digitizer and visualized with the pClamp 11 software package. For sEPSC and intrinsic cellular property recordings, patch pipettes having a tip resistance of 3-5 MΩ contained (in mM): 126 K-Gluconate, 4 KCl, 10 HEPES 4 MgATP, 0.3 Na_2_GTP, 10 PO-Creatine, 0.3% (w/v) Biocytin (pH = 7.25 with KOH, 285 mOsm/kg). For sEPSC recordings, pyramidal neurons were held at −70 mV in voltage-clamp mode. To record intrinsic cellular properties in current-clamp mode, pipette capacitance of 2.0 pF was corrected and series resistance was compensated using the automated bridge balance tool of the Multiclamp commander. IV-curves were generated by injecting 1 s square pulse currents starting at −100 pA and increasing in 10 pA steps until +500 pA injection was reached (sweep duration: 2 seconds). Series resistance was monitored and recordings were discarded if series resistance reached >30 MΩ. For one cell in human cortical slices (Fig. 1, control group), series resistance exceeded 30 MΩ during IV-Curve recording. The respective IV-curve was therefore excluded from further analysis.

### *In vivo* perforant path long-term potentiation

3-month-old male C57BL/6J or synaptopodin-deficient animals (with C57BL/6J genetic background) were kept in a 12 h light/dark cycle (Scantainer) with access to food and water ad libitum. To achieve stable anesthesia, an initial dose of urethane (1.25 g/kg, in sodium chloride solution) was injected subcutaneously (s.c.); 0.1 g/kg supplemental dose as needed. After stable anesthesia was reached, atRA (10 mg/kg in DMSO) or vehicle-only was intraperitoneally injected (blind to experimenter). The mouse was then placed in a stereotactic frame (David Kopf instruments) and local anesthesia with xylocaine (1% s.c. to the scalp) was applied. Cranial access to the brain was established according to coordinates from the mouse brain atlas (Franklin and Paxinos; stimulation electrode: 2.5 mm lateral to the midline, 3.8 mm posterior to bregma; recording electrode: 1.2 mm lateral to the midline, 1.7 mm posterior to bregma). The ground electrode was placed in the neck musculature. Electrophysiological signals were amplified using a Grass P55 A.C. pre-amplifier (Astro-Med, West Warwick) and digitized at 10 kHz sampling rate (Digidata 1440A, Molecular Devices). Extracellular stimulation was performed using a STG1004 stimulator (Multichannel Systems, Reutlingen, Germany). Bipolar stimulation electrode (NE-200, 0.5 mm tip separation, Rhodes Medical Instruments, USA) was lowered 1.5-2.2 mm below the surface of the brain to target the angular bundle of the perforant path. Then the tungsten recording electrode (TM33B01KT, World Precision Instruments, Sarasota, FL, USA) was lowered in 0.1 mm increments while monitoring the waveform in response to 500 μA test pulses until the granule cell layer was reached (1.7-2.2 mm below the surface). The correct placement of the stimulation electrode in the medial portion of the perforant path was verified electrophysiologically by the latency of the population spike, though the activation of some lateral perforant path fibers could not be excluded. Recordings started a minimum of 3 hours after experimental treatment with atRA or vehicle-only. Input-output curve was generated by 30-800 μA current pulses, repeated 3 times at each intensity, 0.1 ms pulse duration, 60 pulses in total at 0.1 Hz. Paired-pulse facilitation was tested by applying two current pulses at increasing inter-pulse intervals (IPIs) from 15 to 100 ms, current intensity set to elicit facilitation of second response, 0.1 ms pulse duration (6 sweeps). Paired-pulse inhibition was tested by two 800 μA current pulses at increasing IPIs from 20 to 1000 ms (0.2 ms pulse duration, 13 sweeps). Recording of perforant path-dentate gyrus (PP/DG)-long-term potentiation (LTP) was performed by applying stimuli with a current intensity set to elicit a 1-2 mV population spike (0.1 Hz, 0.1 ms pulse duration). PP/DG-LTP was induced using a theta-burst stimulation (TBS) protocol comprising three series of six trains with six 400 Hz current pulses at double the baseline intensity and pulse duration (with 200 ms interval between trains and 20 s interval between series). One *Synpo*^*d/d*^ animal in the vehicle-only group was excluded from further analysis, since no rise in the I/O-curve upon increasing stimulus intensities could be detected.

### Immunostaining, post-hoc labeling and light microscopy

Cortical slices were fixed in 4% PFA (prepared from 16% PFA stocks in phosphate buffered saline according to manufacturer’s instruction, Thermo Scientific, #28908) at room temperature and stored at 4°C overnight in the same solution. After fixation, slices were washed in phosphate buffered saline and consecutively incubated for 1 h with 10% (v/v) normal goat serum (NGS) in 0.5% (v/v) Triton X-100 containing PBS to reduce unspecific staining and increase antibody penetration. Subsequently slices were incubated in 10% (v/v) NGS, 0.1% (v/v) Triton X-100 containing PBS at 4°C overnight with rabbit anti-synaptopodin (Synaptic Systems, #163002; 1:1000 dilution) or rabbit anti-NeuN (Abcam, #ab104225, 1:500 dilution) antibodies. Sections were washed in PBS and incubated with suitable goat anti-rabbit AlexaFluor 488- (Invitrogen, #A-11034) or goat anti-rabbit AlexaFluor 555^+^-labeled secondary antibodies (Invitrogen, #A-32732; 1:1000 dilution in 10% (v/v) NGS, 0.1% (v/v) Triton X-100 containing PBS) at 4°C overnight respectively. For post-hoc-visualization of patched pyramidal cells, streptavidin-AlexaFluor 488 (Invitrogen, # S32354; 1:1000 dilution) was added during the secondary antibody incubation. Sections were washed again and incubated for 10 minutes in Sudan Black B (0.1% (w/v) in 70% ethanol) to reduce autofluorescence before incubation with DAPI for 10 minutes (Thermo Scientific, #62248; 1:5000 dilution in PBS) to visualize cytoarchitecture. After final washing, sections were transferred onto glass slides and mounted with fluorescence anti-fading mounting medium (DAKO Fluoromount).

Confocal images were acquired using a Leica SP8 laser-scanning microscope equipped with a 20x multi-immersion (NA 0.75; Leica), a 40x oil-immersion (NA 1.30; Leica) and a 63x oil-immersion objective (NA 1.40; Leica). Image stacks for dendritic spine and synaptopodin cluster analysis were acquired with a 63x objective at 6x optical zoom (resolution: 1024×1024, z-step size: 0.2 μm; at ideal Nyquist rate). Laser intensity and detector gain were set to achieve comparable overall fluorescence intensity throughout stacks between all groups.

### Immunogold labeling of synaptopodin

Human cortical samples were fixed in 0.1% glutaraldehyde and 4% PFA (in phosphate buffer, 0.1 M) for at least 3 hours. 50 μm sections were prepared using a Leica VT1000S vibratome and sections were consecutively incubated with rabbit anti-synaptopodin (Synaptic Systems, #163002; 1:100 dilution in 2% NGS (v/v) containing TBS (Tris-buffered saline; 50 mM)) at 4°C overnight. Sections were washed for 1 hour in TBS (50 mM) and incubated with a suitable secondary goat anti-rabbit antibody (Nanoprobes, #2004; 1.4 nM gold coupled, 1:100 dilution in 2% NGS (v/v) containing TBS (50 mM)) at 4°C overnight. After washing for 30 minutes in TBS (50 mM), sections were postfixed in 1% (w/v) glutaraldehyde containing PBS (25 mM) for 10 minutes. Sections were washed again and a silver intensification (HQ Silver Enhancement Kit, Nanoprobes, #2012) was performed according to the manufacturer’s instruction. Subsequently, slices were incubated with 0.5% osmium tetroxide for 40 minutes, washed in graded ethanol (up to 60% (v/v)) for 10 minutes each and incubated with uranylacetate (1% (w/v) in 70% (v/v) ethanol) for 35 minutes. Slices were then dehydrated in graded ethanol (80%, 90%, 95%, 2x 100% for 10 minutes). Two washing steps in propylenoxide for 5 minutes each were performed before incubation with durcupan/propylenoxide (1:1 for 1 hour) and consecutive transfer to durcupan overnight at room temperature. Slices were embedded in durcupan and ultrathin sectioning (55 nm) was performed using a Leica UC6 Ultracut. Electron micrographs were captured using a Zeiss Leo 906E microscope equipped with a 2kCCD-‘Sharp Eye’ Camera (Tröndle, Moorenweis, Germany) at 10,000x magnification. Acquired images were stored as TIF-files.

### Electron Microscopy

After 6 hours of atRA or vehicle-only treatment, slices were fixed in 4% paraformaldehyde (w/v) and 2.5% glutaraldehyde (w/v; phosphate buffered saline) overnight. After fixation, slices were washed for 4 hours in phosphate buffer (0.1 M). Subsequently, slices were incubated with 1% osmium tetroxide for 45 minutes, washed in graded ethanol (up to 50% (v/v)) for 5 minutes each and incubated with uranylacetate (1% (w/v) in 70% (v/v) ethanol) overnight. Slices were then dehydrated in graded ethanol (80%, 90%, 98% for 5 minutes, 2x 100% for 10 minutes). Subsequently two washing steps in propylenoxide for 10 minutes each were performed before incubation with durcupan/propylenoxide (1:1 for 1 hour) and consecutive transfer to durcupan overnight at room temperature. Slices were embedded in durcupan and ultrathin sectioning (55 nm) was performed using a Leica UC6 Ultracut. Sections were mounted onto copper grids (Plano) and additionally contrasted using Pb-citrate (3 minutes). Electron microscopy was performed at a Philips CM100 microscope equipped with a Gatan camera Orius SC600 at 3900x magnification. Acquired images were saved as TIF-files and analyzed by an investigator blinded to experimental conditions.

### Quantification and statistics

Electrophysiological data were analyzed using Clampfit 11 of the pClamp11 software package (Molecular Devices). sEPSC properties were analyzed by using the automated template search tool for event detection. Immuno-labeled synaptopodin clusters in post-hoc visualized superficial (layer 2/3) pyramidal cells of the human cortex were analyzed in image stacks from 2^nd^ and 3^rd^ order dendritic branches. Synaptopodin clusters that colocalized with dendritic spines (either spine neck or head; Figure 2D) were a target for further analysis. Single plane images of both synaptopodin-positive and -negative clusters were extracted from image stacks, where the spine head cross section area reached its maximum extension and stored as TIF-files. Analysis of spine head cross-sectional area and synaptopodin cluster size was performed manually by a blind investigator using the ImageJ software package (available at http://imagej.nih.gov/ij/). Here, the outer border of synaptopodin clusters and spine heads was marked independently from their overall fluorescence intensity. Data were transferred and stored in an excel-file format. Ultrastructural analysis of the spine apparatus organelles was performed in single plane images of excitatory synapses in the human cortex, where pre- and postsynaptic structures can be readily identified. Cross section area of the spine apparatus organelle was analyzed manually using the ImageJ software package independent of its shape and internal structural organization. *In vivo* perforant path long-term potentiation was analyzed using Clampfit 10.2.

Statistical analysis was performed using the GraphPad Prism 7 software package. Comparison between two groups in one dataset was performed using the Mann-Whitney-U test. U-values were reported for significant differences in the figure legends. To compare more than two groups within one dataset, Kruskal-Wallis test followed by Dunn’s post-hoc correction was used. For statistical evaluation of XY-plots, we used the two-way ANOVA followed by Sidak’s post-hoc correction. In the in-sample analysis (paired experimental design) approach in human cortical slices, we used the Wilcoxon matched-pairs signed rank test. p values smaller 0.05 were considered a significant difference between means. In the text and figures, values represent mean ± standard error of the mean (s.e.m.). * p < 0.05, ** p < 0.01, *** p < 0.001 and not significant differences are indicated by “ns”.

### Graphical illustrations

Confocal image stacks and single plane pictures were stored as TIF-files. Figures were prepared using Photoshop graphics software (Adobe, San Jose, CA, USA). Image brightness and contrast were adjusted.

## REFERENCES

Al Tanoury, Z., Piskunov, A., and Rochette-Egly, C. (2013). Vitamin A and retinoid signaling: genomic and nongenomic effects. J Lipid Res 54, 1761–1775.

Aoto, J., Nam, C.I., Poon, M.M., Ting, P., and Chen, L. (2008). Synaptic signaling by all-trans retinoic acid in homeostatic synaptic plasticity. Neuron 60, 308–320.

Arendt, K.L., Zhang, Y., Jurado, S., Malenka, R.C., Sudhof, T.C., and Chen, L. (2015a). Retinoic Acid and LTP Recruit Postsynaptic AMPA Receptors Using Distinct SNARE-Dependent Mechanisms. Neuron 86, 442–456.

Arendt, K.L., Zhang, Z., Ganesan, S., Hintze, M., Shin, M.M., Tang, Y., Cho, A., Graef, I.A., and Chen, L. (2015b). Calcineurin mediates homeostatic synaptic plasticity by regulating retinoic acid synthesis. Proc Natl Acad Sci U S A 112, E5744–5752.

Biever, A., Glock, C., Tushev, G., Ciirdaeva, E., Dalmay, T., Langer, J.D., and Schuman, E.M. (2020). Monosomes actively translate synaptic mRNAs in neuronal processes. Science 367.

Bosch, M., and Hayashi, Y. (2012). Structural plasticity of dendritic spines. Curr Opin Neurobiol 22, 383–388.

Bremner, J.D., Shearer, K.D., and McCaffery, P.J. (2012). Retinoic acid and affective disorders: the evidence for an association. J Clin Psychiatry 73, 37–50.

Chen, L., Lau, A.G., and Sarti, F. (2014). Synaptic retinoic acid signaling and homeostatic synaptic plasticity. Neuropharmacology 78, 3–12.

Citri, A., and Malenka, R.C. (2008). Synaptic plasticity: multiple forms, functions, and mechanisms. Neuropsychopharmacology 33, 18–41.

Deller, T., Korte, M., Chabanis, S., Drakew, A., Schwegler, H., Stefani, G.G., Zuniga, A., Schwarz, K., Bonhoeffer, T., Zeller, R., et al. (2003). Synaptopodin-deficient mice lack a spine apparatus and show deficits in synaptic plasticity. Proc Natl Acad Sci U S A 100, 10494–10499.

Ding, Y., Qiao, A., Wang, Z., Goodwin, J.S., Lee, E.S., Block, M.L., Allsbrook, M., McDonald, M.P., and Fan, G.H. (2008). Retinoic acid attenuates beta-amyloid deposition and rescues memory deficits in an Alzheimer’s disease transgenic mouse model. J Neurosci 28, 11622–11634.

Dobrotkova, V., Chlapek, P., Mazanek, P., Sterba, J., and Veselska, R. (2018). Traffic lights for retinoids in oncology: molecular markers of retinoid resistance and sensitivity and their use in the management of cancer differentiation therapy. BMC Cancer 18, 1059.

Drager, U.C. (2006). Retinoic acid signaling in the functioning brain. Sci STKE 2006, pe10.

Endres, K., Fahrenholz, F., Lotz, J., Hiemke, C., Teipel, S., Lieb, K., Tuscher, O., and Fellgiebel, A. (2014). Increased CSF APPs-alpha levels in patients with Alzheimer disease treated with acitretin. Neurology 83, 1930–1935.

Gidon, A., Zolnik, T.A., Fidzinski, P., Bolduan, F., Papoutsi, A., Poirazi, P., Holtkamp, M., Vida, I., and Larkum, M.E. (2020). Dendritic action potentials and computation in human layer 2/3 cortical neurons. Science 367, 83–87.

Groth, R.D., and Tsien, R.W. (2008). A role for retinoic acid in homeostatic plasticity. Neuron 60, 192–194.

Hafner, A.S., Donlin-Asp, P.G., Leitch, B., Herzog, E., and Schuman, E.M. (2019). Local protein synthesis is a ubiquitous feature of neuronal pre- and postsynaptic compartments. Science 364.

Ho, V.M., Lee, J.A., and Martin, K.C. (2011). The cell biology of synaptic plasticity. Science 334, 623–628.

Holbro, N., Grunditz, A., and Oertner, T.G. (2009). Differential distribution of endoplasmic reticulum controls metabotropic signaling and plasticity at hippocampal synapses. Proc Natl Acad Sci U S A 106, 15055–15060.

Hu, J., Liu, Y.F., Wu, C.F., Xu, F., Shen, Z.X., Zhu, Y.M., Li, J.M., Tang, W., Zhao, W.L., Wu, W., et al. (2009). Long-term efficacy and safety of all-trans retinoic acid/arsenic trioxide-based therapy in newly diagnosed acute promyelocytic leukemia. Proc Natl Acad Sci U S A 106, 3342–3347.

Koryakina, A., Aeberhard, J., Kiefer, S., Hamburger, M., and Kuenzi, P. (2009). Regulation of secretases by all-trans-retinoic acid. FEBS J 276, 2645–2655.

Kulik, Y.D., Watson, D.J., Cao, G., Kuwajima, M., and Harris, K.M. (2019). Structural plasticity of dendritic secretory compartments during LTP-induced synaptogenesis. Elife 8.

Luo, T., Wagner, E., Crandall, J.E., and Drager, U.C. (2004). A retinoic-acid critical period in the early postnatal mouse brain. Biol Psychiatry 56, 971–980.

Maggio, N., and Vlachos, A. (2014). Synaptic plasticity at the interface of health and disease: New insights on the role of endoplasmic reticulum intracellular calcium stores. Neuroscience 281, 135–146.

Maghsoodi, B., Poon, M.M., Nam, C.I., Aoto, J., Ting, P., and Chen, L. (2008). Retinoic acid regulates RARalpha-mediated control of translation in dendritic RNA granules during homeostatic synaptic plasticity. Proc Natl Acad Sci U S A 105, 16015–16020.

Mansvelder, H.D., Verhoog, M.B., and Goriounova, N.A. (2019). Synaptic plasticity in human cortical circuits: cellular mechanisms of learning and memory in the human brain? Curr Opin Neurobiol 54, 186–193.

Matsuzaki, M., Honkura, N., Ellis-Davies, G.C., and Kasai, H. (2004). Structural basis of long-term potentiation in single dendritic spines. Nature 429, 761–766.

Mundel, P., Heid, H.W., Mundel, T.M., Kruger, M., Reiser, J., and Kriz, W. (1997). Synaptopodin: an actin-associated protein in telencephalic dendrites and renal podocytes. J Cell Biol 139, 193–204.

Poon, M.M., and Chen, L. (2008). Retinoic acid-gated sequence-specific translational control by RARalpha. Proc Natl Acad Sci U S A 105, 20303–20308.

Rataj-Baniowska, M., Niewiadomska-Cimicka, A., Paschaki, M., Szyszka-Niagolov, M., Carramolino, L., Torres, M., Dolle, P., and Krezel, W. (2015). Retinoic Acid Receptor beta Controls Development of Striatonigral Projection Neurons through FGF-Dependent and Meis1-Dependent Mechanisms. J Neurosci 35, 14467–14475.

Reddy, P.H., Mani, G., Park, B.S., Jacques, J., Murdoch, G., Whetsell, W., Jr., Kaye, J., and Manczak, M. (2005). Differential loss of synaptic proteins in Alzheimer’s disease: implications for synaptic dysfunction. J Alzheimers Dis 7, 103–117; discussion 173-180.

Segal, M., Vlachos, A., and Korkotian, E. (2010). The spine apparatus, synaptopodin, and dendritic spine plasticity. Neuroscientist 16, 125–131.

Shearer, K.D., Stoney, P.N., Morgan, P.J., and McCaffery, P.J. (2012). A vitamin for the brain. Trends Neurosci 35, 733–741.

Spacek, J. (1985). Three-dimensional analysis of dendritic spines. II. Spine apparatus and other cytoplasmic components. Anat Embryol (Berl) 171, 235–243.

Sutton, M.A., and Schuman, E.M. (2006). Dendritic protein synthesis, synaptic plasticity, and memory. Cell 127, 49–58.

Ting, J.T., Lee, B.R., Chong, P., Soler-Llavina, G., Cobbs, C., Koch, C., Zeng, H., and Lein, E. (2018). Preparation of Acute Brain Slices Using an Optimized N-Methyl-D-glucamine Protective Recovery Method. J Vis Exp.

Vlachos, A., Ikenberg, B., Lenz, M., Becker, D., Reifenberg, K., Bas-Orth, C., and Deller, T. (2013). Synaptopodin regulates denervation-induced homeostatic synaptic plasticity. Proc Natl Acad Sci U S A 110, 8242–8247.

Vlachos, A., Korkotian, E., Schonfeld, E., Copanaki, E., Deller, T., and Segal, M. (2009). Synaptopodin regulates plasticity of dendritic spines in hippocampal neurons. J Neurosci 29, 1017–1033.

Wingo, A.P., Dammer, E.B., Breen, M.S., Logsdon, B.A., Duong, D.M., Troncosco, J.C., Thambisetty, M., Beach, T.G., Serrano, G.E., Reiman, E.M., et al. (2019). Large-scale proteomic analysis of human brain identifies proteins associated with cognitive trajectory in advanced age. Nat Commun 10, 1619.

Zhang, Z., Marro, S.G., Zhang, Y., Arendt, K.L., Patzke, C., Zhou, B., Fair, T., Yang, N., Sudhof, T.C., Wernig, M., et al. (2018). The fragile X mutation impairs homeostatic plasticity in human neurons by blocking synaptic retinoic acid signaling. Sci Transl Med 10.

