## Supplementary Material for "All-Trans Retinoic Acid induces synaptic plasticity in human cortical neurons"

### Supplemental Table S1: Cortical resection samples supplementary information

| sample | age [y] | sex | pathology | region | site |
|---|---|---|---|---|---|
| **1** | 47 | m | epilepsy | temporal | right |
| **2** | 59 | m | tumor | frontal | right |
| **3** | 56 | m | tumor | frontal | left |
| **4** | 70 | f | tumor | temporal | left |
| **5*** | 55 | m | tumor | temporal | right |
| **6** | 55 | m | tumor | frontal | right |
| **7** | 52 | m | tumor | frontal | left |
| **summary** | Ø = 56.3 years | m:6 / f:1 | e:1 / t:6 | temp.:3 / front.:4 | r:4 / l:3 |

*Sample 5 has been used for anisomycin experiments only.

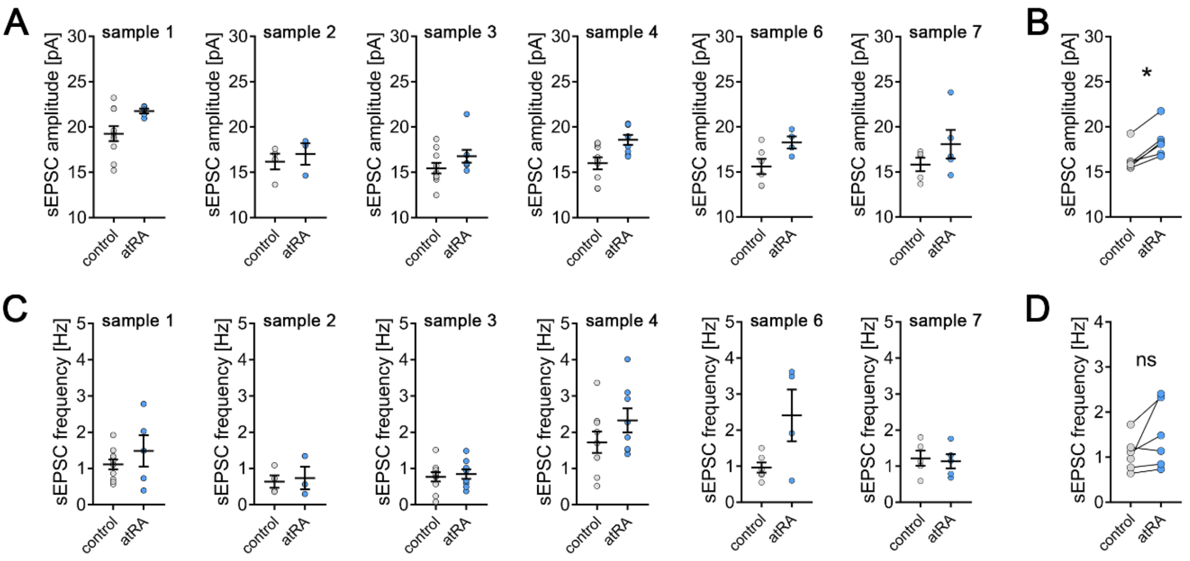


**Figure S1. All-trans retinoic acid (atRA) induces excitatory synaptic strengthening in human superficial (layer 2/3) pyramidal neurons – an in-sample control analysis.**

**(A)** A total amount of 7 human neocortical samples were collected for this study, while cortical slices from 6 samples were included in this dataset. In each sample acute cortical slices were randomly assigned to the atRA- or vehicle-only-treated group. In every sample an increase in sEPSC amplitude can be observed ($n_{control}$ = 43 cells, $n_{atRA}$ = 33 cells in 6 samples each; sample 1: $n_{control}$ = 10 cells, $n_{atRA}$ = 5 cells (one cell excluded from further analysis with 38.4 pA/6.5 Hz sEPSC amplitude/frequency respectively); sample 2: $n_{control}$ = 4 cells, $n_{atRA}$ = 3 cells; sample 3: $n_{control}$ = 10 cells, $n_{atRA}$ = 8 cells; sample 4: $n_{control}$ = 9 cells, $n_{atRA}$ = 8 cells; sample 6: $n_{control}$ = 6 cells, $n_{atRA}$ = 4 cells; sample 7: $n_{control}$ = 5 cells, $n_{atRA}$ = 5 cells). **(B)** Paired statistical analysis of mean sEPSC amplitude of atRA-treated slices and their respective in-sample vehicle-only-treated slices reveals a significant increase in sEPSC amplitude following atRA treatment (n = 6 samples; Wilcoxon matched-pairs signed rank test). **(C, D)** Analysis of sEPSC frequency with a paired statistical approach does not reveal a significant increase in sEPSC frequency upon atRA treatment.

Individual data points are indicated by gray or blue dots respectively. Values represent mean ± s.e.m. (ns, non-significant difference, * $p < 0.05$).

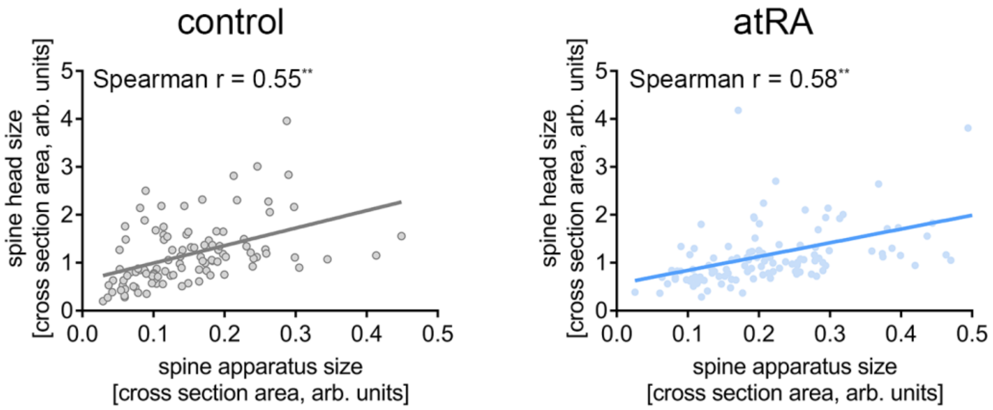


**Figure S2. Ultrastructural analysis of excitatory synapses reveals a positive correlation between sizes of the spine head and the spine apparatus organelle.** XY-plots of paired spine head and spine apparatus organelle size (cross section area) were created for both vehicle-only- and atRA-treated human neocortical slices. A positive correlation can be detected that is preserved following atRA treatment ($n_{control}$ = 103, $n_{atRA}$ = 114; linear regression fit, Spearman $r_{control}$ = 0.55** and $r_{atRA}$ = 0.58**).

Individual data points are indicated by gray or blue dots respectively (** $p < 0.01$).
